# *Rhodococcus equi* Escapes Macrophage Autophagy: VapA Inhibits ATP6V0d1-Mediated Lysosomal Degradation

**DOI:** 10.1101/2023.10.26.564126

**Authors:** Haixia Luo, Tingting Feng, Zhaokun Xu

**Author notes:** Correspondence: Haixia Luo.

## Abstract

*Rhodococcus equi* is a gram-positive actinomycetales bacterial, which cause a pneumonia in foals and in immunocompromised humans. Virulence associated protein A (VapA) encoded from virulence plasmid is essential for intracellular proliferation in macrophage. It has reported that VapA participates in exclusion of proton-pumping vacuolar-ATPase complex from phagosome and causes membrane permeabilization, thus contributing to a pH-neutral phagosome lumen. However, there have been no reports about VapA-induced autophagy or its mechanism. In this study, we sought to determine the role of VapA in macrophage autophagy using western blotting, immunofluorescence, pull-down, Co-IP, LS/MS, pH detection LysoTracker and CFU analysis. We show that VapA inhibiting the macrophage autophagy by detecting the expression of autophagy related protein LC3II and P62 and autophagy flux in VapA treated macrophage. Furthermore, we using pull-down and MS analysis to selected ATP6V0D1 could interact with VapA, Co-IP was used to confirm its interaction. Furthermore, VapA treatment in ATP6V0D1 overexpressed cells weaken the lyso Tracker red stain comparing with ATP6V0D1 overexpressed J774A.1 cells. In addition, the expression of ATP6V0D1 significantly increased in J774A.1 post infected with vapA deletion strain, *R. equi 103*^*+*^/ΔVapA by comparing with wild type *R. equi 103*^*+*^ post-infection. And raise of raise of ATP6V0D1 actually reduce *R. equi 103*^+^ intracellular multiplication. These results further suggesting a decrease of ATP6V0D1 were caused the exit of VapA in *R. equi*. Taken those date together, we reported that VapA mediated lysosome acidification by inhibiting the expression of ATP6V0D1, which contributing the *R. equi* escape macrophage autophagy.

## Introduction

*Rhodococcus equi* (*R. equi*) is a gram-positive bacterial, which infects foals between 3-5 months old and cause a severe pyogranulomatous pneumonia [1]. It can also infect patients with immunodeficiency disease [2]. *R. equi* is phagocytosed and resides in an early phagosome compartment, which fail to undergo later maturation stages thereby resulting in an *R. equi* containing vacuole (RCV). Formation of RCV results in bacterial multiplication and killing of the host macrophage by necrosis with the net effect of *R. equi* escaping macrophage bactericidal activity [3, 4]. A plasmid encoding Virulence-associated-protein A (VapA) is essential for *R. equi* intracellular survival in macrophages [5]. VapA is cell surface protein and produced in 37°C. There are several VapA homologues proteins presented in pathogenic Ireland [6, 7]. Vap homologues were found from diverse species, including *Xenorhadus bovienii, Halomonas titaniciae, Lacinutrix, Escherichia coli* and *Clostridium* [8]. The molecular function of VapA and VapA-like protein remained elusive in spite of the availability of crystal structures of Vap family members [8].

Autophagy is a highly conserved, intracellular degradative process, which involves the engulfment of a protein of cell cytoplasm and organelles in double membrane vesicle known as autophagsome, which then fuses with the lysosome for degradation [9]. Proton-pumping vacuolar ATPase (V-ATPase) is a large multiple subunit complex from the phagosome, which consist of a cytosolic V_1_ domain with 8 subunits that mediates ATP hydrolysis, and an integral membrane V_0_ domain with 6 subunits in yeast and additional AC45 protein in more complex eukaryotes that has proton pump activity. The activities of lysosomal hydrolases require an acidic environment, which is achieved by the proton pump activity of V-ATPase. It has reported that VapA disrupts autophagosomes and/or endosomes fuse with lysosomes to form an autolysosome or an endolysosome, but the molecular basis of this observation or VapA induced autophagy remained undetermined [10, 11]. Following up on this observation, we show here that VapA could inhibit the late autophagy. We further address that VapA mediate ATP6V0D1 to destroy the lysosome acidification, which contributing the *R. equi* escape macrophage autophagy.

## Materials and Methods

### Antibodies and reagents

Anti-VapA was produced by ABclonal Technology Co.,Ltd (Wuhan, China). Anti-LC3, anti-p62, anti-GAPDH and Goat anti-rabbit and anti-mouse IgG-horseradish peroxidase (HRP) were purchased form Proteintech (Shanghai, China). Anti-ATP6V0D1 was purchased form Abcam (Cambridge, UK). Fetal bovine serum (FBS) and Dulbecco’s modified Eagle’s medium (DMED) and Phosphate buffered saline (PBS) were purchased from Biological Industries (China marketing department, Israel). Protein markers and Lipofectamine RNA iMAX Transfection Reagent and Lipofectamine 2000 Transfection Reagent and His pull-down kit and Secondary fluorescent antibodies were purchased form Thermo Fisher scientific (Massachusetts, USA). Lyso-Tracker Red and Hoechst 33342 live cell staining and RIPA buffer and DAPI dye and BCA Protein Assay Kit were purchased form Beyotime Biotechnology (Shanghai, China). Brain heart infusion and LB solid medium and Bovine albumin (BSA) were purchased form Solarbio (Beijing, China). Pull down kit was purchased form Beaver(Suzhou, China).

### Strains and Cell culture

Competent cell *DH5α* strains were purchased from TransGen Biotech (Beijing, China). *R. equi 103*^+^, plasmid-less and avirulent strain *R. equi 103*^-^(*R. equi 103*^-^) and *vapA* deletion strain (*R. equi103*^+^/ △ *vapA*) were kindly provided by Wim G Meijer (University College Dublin, Ireland). *R. equi* were grown on brain heart infusion (BHI) at 37°C, 200 rpm on shaker. The murine macrophage cell line J774A.1 was obtained from the American Type Culture Collection. J774A.1 were cultivated with DMEM supplement with 10% FBS at 37°C, 5% CO_2_.

### Immunoblotting

J774A.1 were seeded at 2×10^6^ cells per well on 6-well plates and incubated for overnight. Purified VapA protein (100μg/ml) treated J774A.1 cell for 24h or BafilomycinA (50mM) for 2h. Collected J774A.1 cells were washed by PBS for three tines and lysed with RIPA buffer for 5 min at 4°C. Then centrifuged at 13,000 x g for 5 min. Protein were quantified using BCA Protein Assay Kit. Proteins were transferred to nitrocellulose membranes used the Western Blot system (Bio-Rad) after SDS-PAGE. Membranes were blocked in 5% (w/v) semi-skimmed milk for 1 h at room temperature, and then incubated with overnight at 4°C with primary antibodies, Anti-LC3 (1:1000), anti-p62 (1:1000), Anti-ATP6V0D1 (1:1000), anti-GAPDH (1:1000) diluted in block buffer. After then, the membrane was incubated with IgG-horseradish peroxidase (HRP) second antibody (1:10000) for 1 h at room temperature and then proteins were visualized with enhance chemiluminescence (ECL) kit purchased from GE Healthcare (Chicago, USA). And detection reagent and analyzed using GE Amersham Imager 600.

### Immunofluorescence microscopy detect LC3 and P62

1×10^5^ J774A.1 cells were seeded on glassslides in 12 well plagtes for overnight. 100μg/ml purified VapA protein was added per well for 24 h, 50μg/ml Bafilomycin A was added per well for 2 h. The cells were washed twice with PBS, permeabilized with 0.05 % (w/v) Triton X-100 and blocked in 5% (w/v) BSA for 30 min at room temperature. Primary antibody Anti-LC3B (1:1000) and anti-P62 (1:1000) were incubated overnight at 4 °C. Then washed three times for 5min with blocking buffer and secondary fluorescent antibodies were added for 1 h at room temperature. Fluorescent images were obtained using confocal microscope (Wezlar, Germany).

### Autophagy flux

Tandem-fluores-cent-tagged LC3 reporter(mRFP-GFP-LC3) infection J774A.1 for 48hr, then treatment with recombinant VapA(100μg/ml) protein 24hr. Cells were washed PBS (2×5 min), permeabilized with 0.05% (w/v) Triton X-100 for 30 min and block with Antifade Solution. Samples were recorded through a laser scanning confocal microscope (Wezlar, Germany).

### Pull-down and LS/MS analysis

J774A.1 were collected and washed by PBS for three times and lysed with Lysis buffer and prepare recombinant VapA protein for 200μg. Pipette 50μL of the slurry into each labeled spin column and used washing solution wash for 5 times. Recombinant VapA protein and slurry incubate at 4°C for 2hr with gentle rocking motion on a rotating platform. Used wash solution washed for 5 times. Add up to 800μL of cell Lysis solution (600μg) and incubate at 4°C for 2hr with gentle rocking motion on a rotating platform. Used wash solution washed for 5 times and used elution Buffer incubate spin column for 10 minutes with gentle rocking. Used Triple TOF 5600(SCIEX, USA) LC-MS system to mass spectrometry detection. The data were retrieved in the mouse proteome database through maxQuant (v1.6.2.10) for further analysis

### Immunoprecipitation

100μg/ml VapA treated J774A.1 cells we treated with protein for 24 h. Cell were washed by PBS for three times and lysed with RIPA buffer for 5 mins at 4°C, centrifuged at 13,000xg for 5 min at 4°C to collect the supernatant. The supernatant was incubated with VapA primary antibody (1:200) at 4°C for overnight then followed by incubation with protein G magnetic bead for 2h at 4°C with rotate mixture. After five washes with lysis buffer, the immunocomplexes bound beads were 99°C treatment for 10 mins then analyzed by SDS-PAGE and immunoblotting with distinctive antibodies.

### transient transfection of siRNA and eukaryotic expression vector

*Atp6v0d1* siRNA were transfected into J774A.1 using Lipofectamine RNA iMAX transfection reagent. The sequences are as follows: Atp6v0d1 siRNA forward:(CAUCGAGAUAAUCCGAAAUTT), Atp6v0d1 siRNA reverse:(AUUUCGGAUUAUCUCGAUGTT). Atp6v0d1 eukaryotic expression vector was gene synthesis form General Biotechnology Co., Ltd (Anhui, China). Control empty vector and pcDNA3.1-*Atp6v0d1* transfected into J774A.1 using Lipofectamine 2000 transfection reagent. Used SDS-PAGE and Western blotting to detection the expression.

### LysoTracker

J774A.1 were seeded at 1×10^5^ cells per well on glass slides and cultured for overnight. After transient transfection of siRNA and eukaryotic expression vector, treatment with 100 μg/ml recombinant VapA protein and *R.equi* strains, the cells were incubated with Lyso-tracker Red for 1h at 37°, 5% CO2. Hoechst 33342 living cell staining solution was incubated at 37°, 5% CO2 for 10 min. Samples were recorded through fluorescence microscope(Wezlar, Germany).

### CFU

J774A.1 were seeded at 2×10^6^ cells per well on 6-well plates and cultured for overnight. *R. equi* was fostered in BHI culture medium with shaking at 37°C to reach an OD600 of 0.6-0.8. And washed with PBS for twice times. Bacteria were taken to infect J774A.1 at MOI (multiplicity of infection) of 10. Bacteria were centrifuged at 160g for 3 min and incubated for 1hr at 37°C to allow the cell phagocytosis of bacteria. The cell was rinsed three times with DMEM to remove unbound bacteria, and the media replaced with DMEM containing 10% FBS and 5μg/ml vancomycin to kill extracellular bacteria. Used sterile water containing the sample and place it at room temperature for 5 min to make the cells absorbed water and busted to release the intracellular bacteria. Doubling dilution and culture with LB solid medium at 37°C. After 36 h record the data for subsequent experimental analysis.

### Statistical analysis and image preparation

Statistical analysis applied GraphPad Prism 7 software and data from three groups were compared by one-way analysis of variance (ANOVA) followed. When p-values lower than 0.05 were considered significant.

## Results

### VapA inhibits macrophage autophagy

Autophagy related protein LC3 and P62 are key proteins associated with autophagy. Transformation of LC3 from the cytosolic type (LC3-I) to autophagosomal-associated type (LC3-II) is commonly used as a specific marker for autophagosome formation [12]. P62 is autophagic adaptor, which binds to ubiquitinated cargo via the UBA domain and LC3 Via its LC3 interaction region (LIR). This step helps to sequestration of cargo autophagosomes. P62 is degraded by autophagy and hence decreased autophagic flux leads to accumulation of this protein. To investigate the role of VapA on macrophage autophagy, we measure the LC3-II and P62 level in J774A.1 cells treated with VapA in doze and time dependent manner. We treated J774A.1 cell with 5-200 μg/ml VapA for 24h. LC3-II and P62 were increased significantly in a dose-dependent manner, where as it was observed that 100 μM was the most effective concentration (Fig.1A). Then, 100 μg/ml VapA was used to treat the J774A.1 cells for 3 h, 6 h, 12 h, 24 h and 48 h. LC3-II and P62 were increased significantly comparing with untreated cell (control). We observed 24 h treatment of VapA show the most accumulation of LC3-II and P62 (Fig.1B). Therefore, we are using 100 μg/ml and 24 h treatment of VapA for the further experiment.

**Figure 1.**
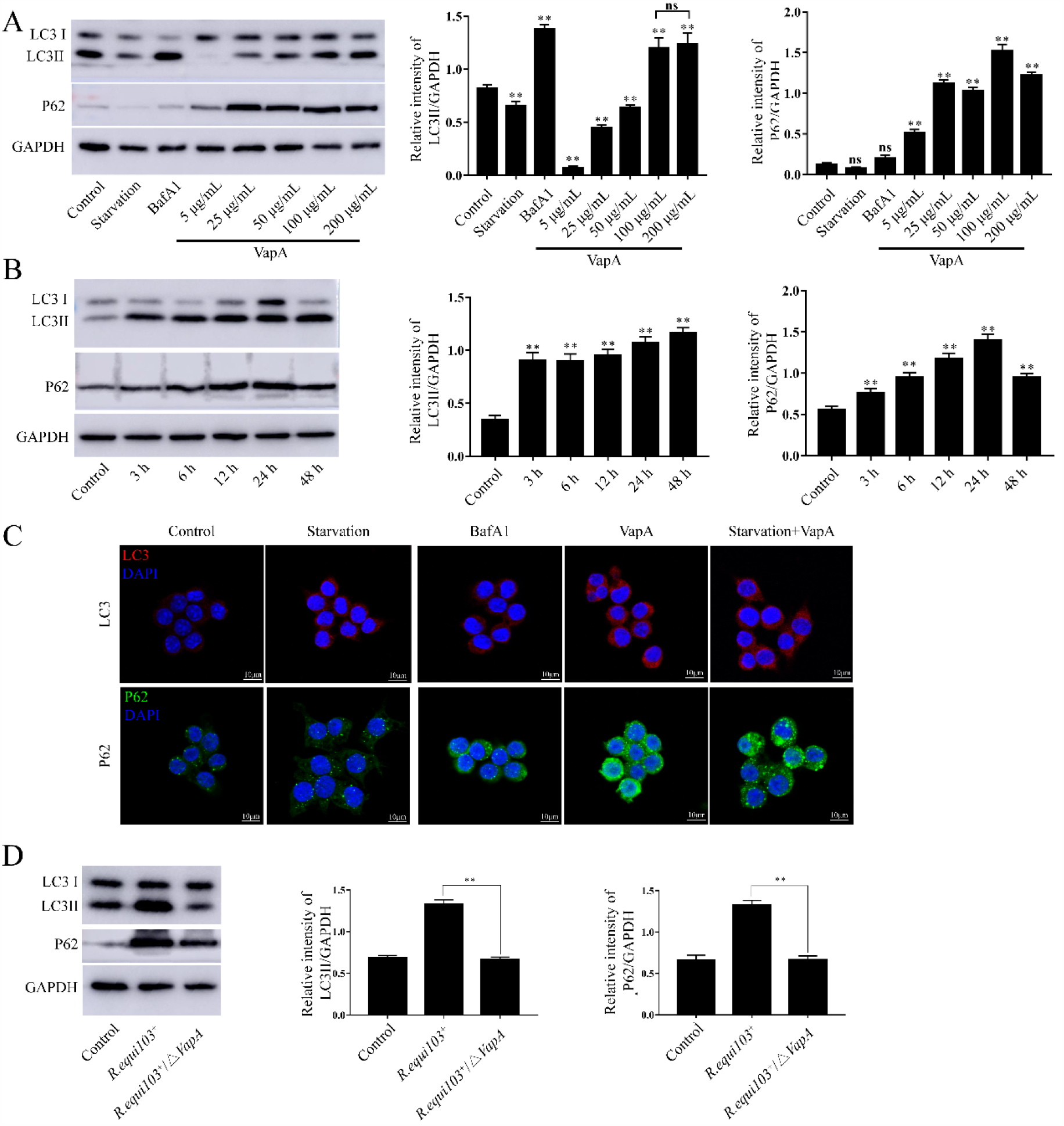
VapA can inhibit autophagy in J774A.1 cells. (A) Expression of LC3 and P62 el in J774A.1 cells treated with 5-200 μg/ml (B) Expression of LC3 and P62 el in J774A.1 with treatment of VapA for 3 h, 6 h, 12 h, 24 h and 48 h. (C) Autophagosome accumulation revealed by LC3-II and P62 accumulation in J774A.1 cells with VapA and/or starvation treatment. Scale bars 25 μm. (D) Expression of LC3-II and P62 were measured by western blotting in J774A.1 cells postinfection with virulence *R.equi103+* (MOI of 10) and *vapA* deletion strain *R.equi103+/*Δ*vapA* (MOI of 10) for 24h. The data represent the mean ± SD from three independent experiments performed in triplicate.

With LC3-II increasing and P62 accumulating in VapA treatment cell from above results, which indicate that VapA might block in autophagic flux. Starvation is an autophagy inducer. Bafilomycin A1 (BafA1) block the fusion of autophagosome with lysosomes. To further test VapA mediated macrophage autophagy, we compared the levels of LC3-II and P62 in J774A.1 cells were then stimulated with starvation(FBS free treatment for 2h), BafA1 (50mM), VapA (100 μg/ml), VapA (100 μg/ml) pus starvation (starvation+VapA) treatment, and confocal microscopy analysis LC3-II increased, VapA and VapA+starvation caused an increase of LC3-II were similar to BafA1. VapA and VapA+starvation inhibited p62 degradation compared to Satravation or BafA1 treated J774A.1 cells. The combined treatment of starvation and VapA cause further accumulation of p62, suggesting that VapA can inhibit rather than induce autophagic flux (Fig1C).

*R.equi* wild type stain, *R.equi103*^*+*^ and *vapA* deletion strain, *R.equi103*^*+*^*/*Δ*vapA* were used to determine the relationship between VapA and the accumulation of LC3 and P62 in J774A.1. As shown in figure 5D, comping with uninfected cell, the LC3-II and P62 were significantly increased in J774.1 post-infected with *R.equi103*^*+*^. While, the expression of LC3-II and P62 were significantly decreased in *R.equi103*^*+*^*/*Δ*vapA* infected macrophage comping with *R.equi103*^*+*^ post-infection. From above results, we showed that VapA inhibits the macrophage autophagy.

To validate these observations and further dissect the step of autophagic flux affect by VapA, we employed an adenovirus mRFP-GFP-LC3 to measure the autophagic flux, while RFP-positive puncta (RFP^+^ GFP^+^) detect autophagosomes appear yellow, detect autolysosome (the fusion product of autophagosomes with lysosomes) are seen as red pucta (RFP^+^ GFP-) because the green fluorescence of GFP is quenched due the acidic nature of lysosomes. J774A.1 cells transfected with adenovirus mRFP-GFP-LC3, then treated with VapA (100 μg/ml) for 24 h on nutrition or starvation situation. We saw a significant increase in the number of autophagsomes (mRFP^+^/GFP^+^) and autolysosomes (mRFP^+^/GFP^-^) in starvation treated cells, VapA and VapA+starvation treatment comparing with untreated cell (control). While, Comparing with starvation treatment, the autophagsomes and autolysosomes were significant decreased in VapA and VapA+starvation treatment J774A.1. These results t suggested that VapA inhibits autophagic flux most likely happens at the autolysosomes stage.

### VapA inhibit the expression of ATP6V0d1

We next sought to identify the potential autophagy related interacting proteins of VapA, we performed pull-down and LC-MS/MS assay in this study. Purified VapA protein was used as a bait protein to pull-down the relevant protein in *R.equi 103*^*+*^ infected macrophage. As shown in Fig.3A, the significant difference bands were detected between VapA pull-down eluent (Exp) and control rabbit IgG (Control). Then, these two elute samples were subjected to LC-MS/MS analysis. The LC-MS/MS spectra was searched against the mice protein Database using the mascot software, which led to the identification of 258 difference abundance proteins (data not show). Through GO and signal pathway analysis, Seven autophagy related proteins were identified (Table 1). It reported that VapA neutralizes the phagosome lumen by exclusion of vATPase from the phagosome limiting membrane and by permeabilisation for protons [11]. Anti-VapA and anti-ATP6V0d1 antibodies were used to verify the VapA and ATP6V0d1 in pull down elute solution. Both VapA and ATP6V0d1 were detected in Exp elute solution, but not in control (fig.3B). We also performed a Co-IP using IgG beads, VapA and ATP6V0d1 could be found in the immunoprecipitated lysates of VapA treated J774A.1 cells (fig.3C). Those results proved that ATP6V0d1 could interact with VapA.

**Table:** The interaction with with VapA by LC-MS/MS.

**Table: The interaction with with VapA by LC-MS/MS**
| **Name** | **Accession** | **Description** |
| --- | --- | --- |
| Pdia6 | Q922R8 | Protein disulfide-isomerase A6 OS=Mus musculus<br>OX=10090 GN=Pdia6 PE=1 SV=3 |
| Cisd2 | Q9CQB5 | CDGSH iron-sulfur domain-containing protein 2 OS=Mus<br>musculus OX=10090 GN=Cisd2 PE=1 SV=1 |
| Vcp | Q01853 | Transitional endoplasmic reticulum ATPase OS=Mus<br>musculus OX=10090 GN=Vcp PE=1 SV=4 |
| Arpc3 | Q9JM76 | Actin-related protein 2/3 complex subunit 3 OS=Mus<br>musculus OX=10090 GN=Arpc3 PE=1 SV=3 |
| Actr3 | Q99JY9 | Actin-related protein 3 OS=Mus musculus OX=10090<br>GN=Actr3 PE=1 SV=3 |
| Mapk3 | Q63844 | Mitogen-activated protein kinase 3 OS=Mus musculus<br>OX=10090 GN=Mapk3 PE=1 SV=5 |
| Atp6v0d1 | P51863 | V-type proton ATPase subunit d 1 OS=Mus musculus<br>OX=10090 GN=Atp6v0d1 PE=1 SV=2 |

**Figure 2.**
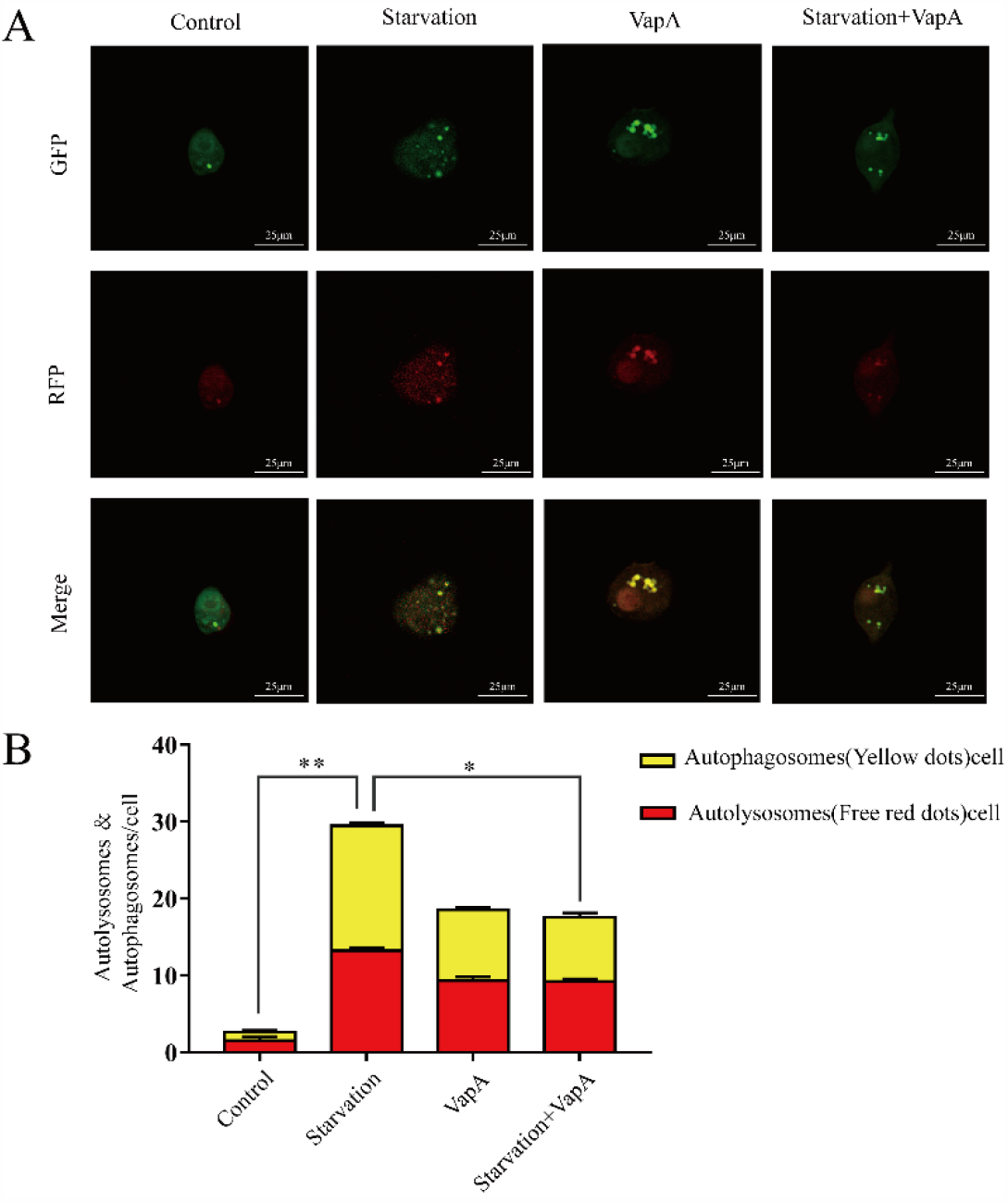
VapA inhibiting autophagic flux in J77A4A.1 cells. J774A.1. J774A.1 cells were transfected with mRFP-GFP-LC3-I/II (red and green) followed by starvation, VapA (100 μg/ml) or VapA+starvation treatment. The amount of autophagosome (yellow) and autolysosome (red) per cell was determined using Impair software. Scale bar 25μm. ^*^,<0.05; ^**^,<0.01 versus control group or starvation.

**Figure 3.**
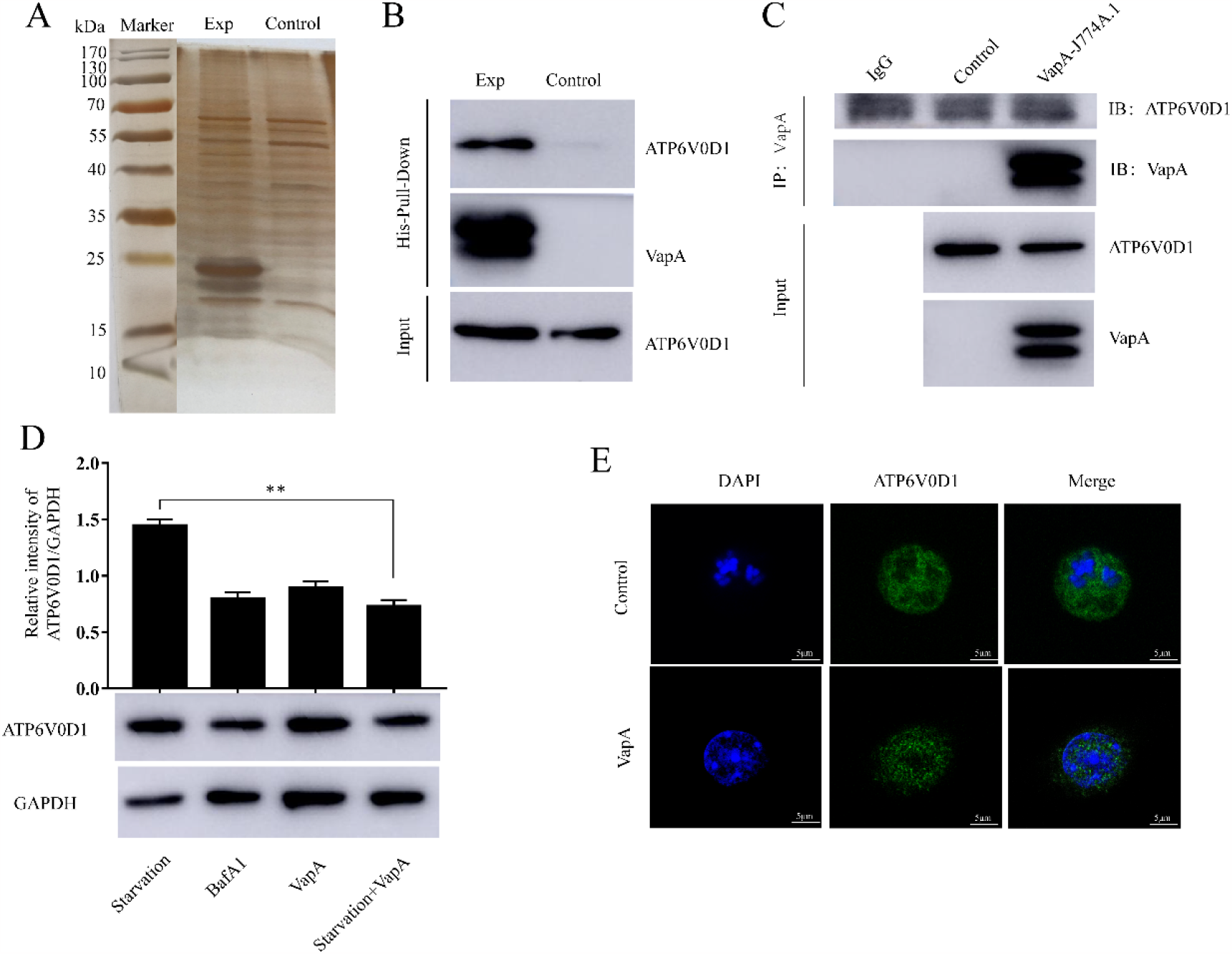
VapA inhibit ATP6V0d1 expression in macrophage. (A) The eluted protein were detected with SDS-PAGE. (B) The eluted proteins were detected with anti-VapA and anti-ATP6V0d1 antibodies. (C) VapA and ATP6V0d1 were preincubated for overnight at 4°C and immunoprecipitated with anti-VapA and anti-ATP6V0d1 antibody. IP, immunoprecitated; IB, immunoblotted. (D and E)Western blot and Immunofluorescence microsopy analysis determined protein levels of ATP6V0d1 in J774A.1 cells with starvation, BafA1 (50mM), VapA (100 μg/ml) and VapA (100 μg/ml) +starvation treatment.

To test the involvement of ATP6V0d1 in VapA induced autophagy, we measure the Atp6v0d1 expression in J774A.1 cells treated with starvation, BafA1 (50mM), VapA (100μg/ml) +starvation(FBS free treatment for 2h). Western blot analysis demonstrated that ATP6V0D1 was significantly decreased in J774A.1 cells following BafA1, VapA and VapA+starvation treatment comparing starvation treatment (Fig 3D). Immunofluorescence was employed to dissect the effect of VapA on the expression of ATP6V0d1. A similar decrease expression of ATP6V0d1 was observed in VapA-treated J77A4.1 cell compring with untreated J774A.1 (control) (fig 3E). These findings suggested that VapA inhibited macrophage autophagy by inhibiting of ATP6V0d1.

### VapA affect endo-lysosomal function by interaction with ATP6V0D1

LysoTracker is a membrane-permeable fluorescent dye that is captured by protonation in compartments with a PH of 6 or lower, such as late endosomes, late phagosomes, lysosomes and autolysosomes [13]. VapA could generate a ph-neutral and hence growth–promoting intracellular niche in macrophage. To investigate whether VapA generated ph-neutral was coursed by the inhibiting of ATP6V0D1, we used LysoTraker deep red, which preferably accumulates in acidic compartments in ATP6V0D1 overexpressed J77A.1 cells by transfected *vapA* eukaryotic expression vector pcDNA3.1-*Atpv60D1*. In treatment with VapA prior to ATP6V0D1 overexpress cells the lyso Tracker red staining had a weaker staining than ATP6V0D1 overexpressed cells (Fig3C). These findings suggested that VapA mediate lysosome acidification by inhibiting the expression of ATP6V0D1.

### R.equi escape macrophage clearance by VapA prevent ATP6V0D1 expression

We reasoned that VapA mediated lysosome acidification by inhibiting ATP6V0d1, and then increasing the ATP6V0D1 by the other means might turn macrophage into susceptible host for aviurlent *R.equi* strain. The virulent strain *R.equi103+* or *vapA* deletion strain (*R.equi 103+/*Δ*VapA*). The levels of ATP6V0D1 were detected in J774A.1 following *R.equi 103+* or *R.equi 103+/*Δ*VapA* infection for 24 h.The expression of ATP6V0D1 increased in J774A.1 post infected with *R.equi 103+/*Δ*VapA* by comparing with *R.equi 103+* post-infection. These results further suggesting a decrease of ATP6V0D1 were caused the exit of VapA in *R.equi*.

To explore the effect of ATP6V0D1 on *R.equi* intracellular survival, the overexpress ATP6V0D1 J774A.1 cells were used in this study. We detected the intracellular proliferation of *R.equi 103+* and *R.equi 103+/*Δ*VapA* at 24 h and 48 h. We noted that the raise of ATPV0D1 actually reduce *R.equi 103+* intracellular multiplication. Together, these data indicate that VapA affected lysosome acidication by inhibiting the expression of ATPV0D1, which play an important role for *R.equi* intracellular proliferation. Taken together, these data suggest ATPV0D1 in some extend is resist virulence R.equi103+ intracellular proliferation by promotion of autophagosome-lysosome.

## 3. Discussion

*R.equi* is a major cause of pneumonia in foals, it also a great challenge to the immunocompromised adult. This disease continues to be a major challenge for clinicians and the equine industry in terms of epidemiological pattern and the therapeutic control due to its complex host-pathogen interaction. Autophagy is a cytoprotective mechanism in which its own cytoplasmic contents are engulfed into autophagosomes and destined for lysosomal degradation. The autophagy mechanism is well studied in various disease states such as cancer, neurodegenerative diseases, aging and immunity and inflammation [14]([1] Deretic V. Autophagy in inflammation, infection, and immunometabolism[J]. Immunity, 2021, 54(3):437-453.). In order to survive and replicate within host cells, some microorganisms have evolved a mechanism to escape degradation by the autophagy machinery. It has been shown that mycobacterial species, including *M.tuberculosis* and *M. marinum* survival in macrophages by preventing both phagolysosome formation and recruitment of LC3 to its phagosome[15] [16]. *R.equi* interferes with phagosomal maturation and suppresses of phagolysosomes[17] [18]. Virulence associated protein A (VapA) encoded from virulence plasmid is a crucial virulence factor for *R.equi* pathogen in macrophage, which play an important role in suppresses of phagolysosomes process [17]. It is reasonable to speculate that VapA might inhibiting the late autophagy to control infection by intracellular *R.equi*. To prove this hypothesis, we present autophagy associated protein LC3-II and P62 is increased in VapA treated cells and virulence strain 103+ post-infected macrophage (Fig.1 and Fig4). In the contrast, the expression of LC3-II and p62 is decreased in macrophage post-infected with vapA deletion strain (Fig 4) and VapA inhibiting starvation induced autophagy by combine the treatment starvation and VapA together (Fig1). It show the evidence that that VapA interfere the autophagy by affect the function of lysosomal.

**Figure 4.**
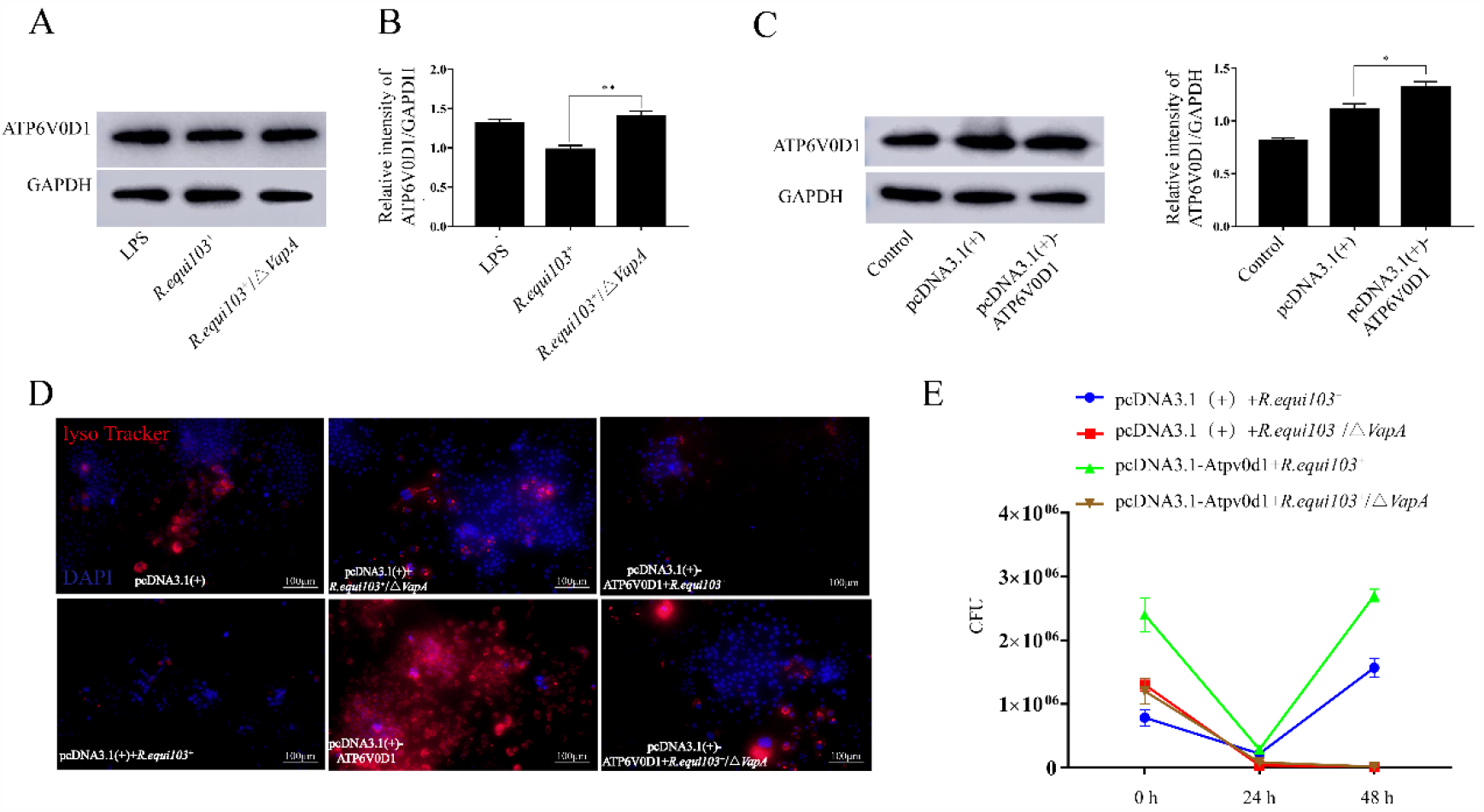
VapA affect endo-lysosomal function by interaction with ATP6V0d1. (A and B) western blot analysis determined protein levels of ATP6V0d1 in J774A.1 cells transfected vapA eukaryotic expression vector pcDNA3.1-Atpv60d1. J774A.1 cell transfected with empty vector pcDNA3.1 used as a control. (B) LysoTracker strain for J774A.1 treated with VapA prior to transfected with pcDNA3.1-Atpv60d1 or pcDNA3.1.

**Fig 5.**
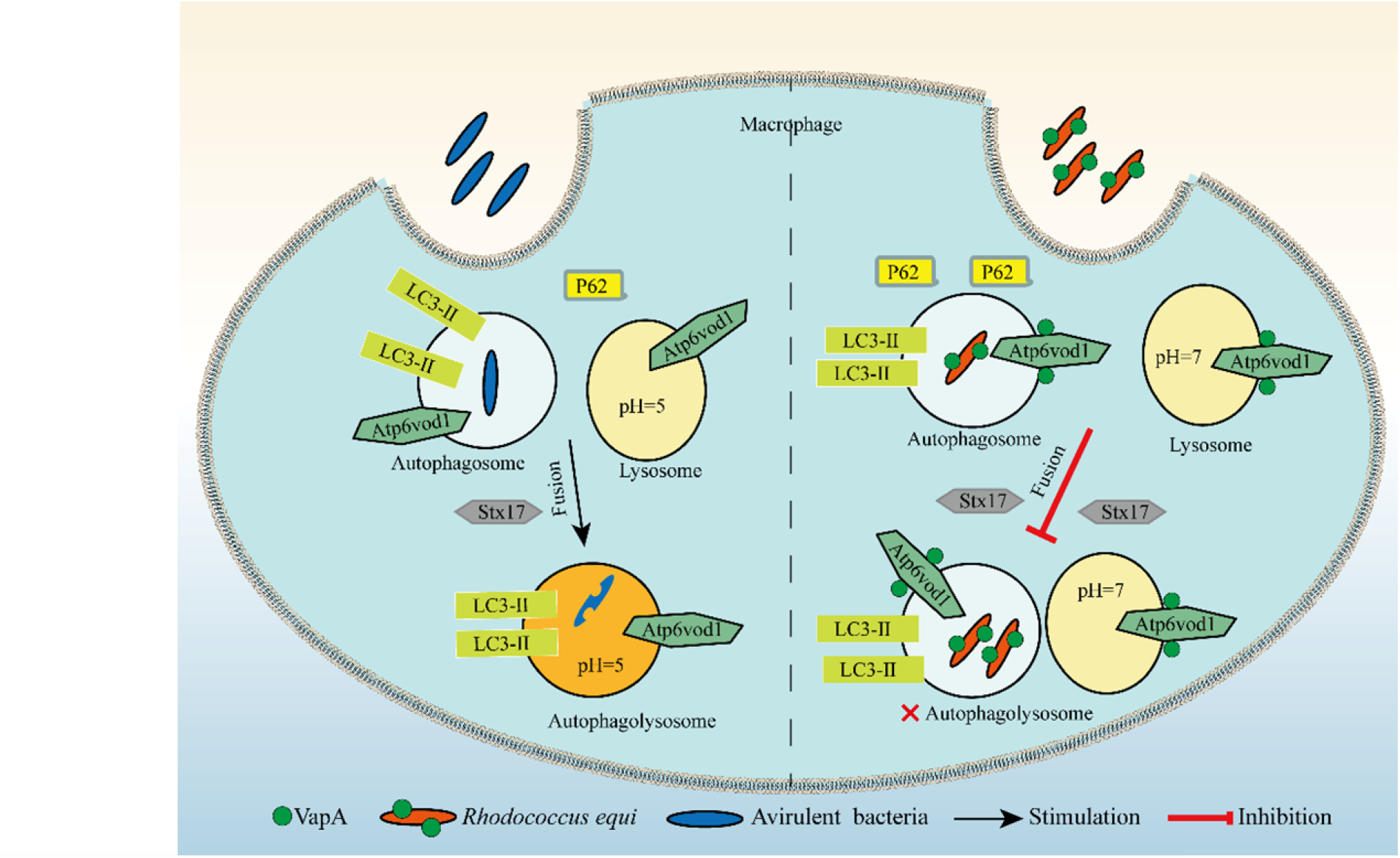
Model of VapA inhibiting Atp6vod1 affect autophagolysosomal in macrophage.

The activities of lysosomal hydrolases require acidic environment, which is achieved by the proton pump activity of V-ATPase[19]. V-ATPase is a large multiple subunit complex consisting of cytosolic V1 domain with 8 subunits that mediates ATP hydrolysis. There are 2 isoforms of ATPV0D subunit, ATPV0D1 and ATPV0D2 (16). ATPV0D1 is ubiquitously expressed, ATPV0D2 has more restricted tissue expression, in osteoclasts and kidney tubules (17-19). We found that ATPV0D1 play a complementary role in VapA inhibited macrophage autophagy. *R.equi* has been shown to inhibit phagolysosomal fusion, it is reasonable to speculate that infection with R.equi could induce the same innate host response as infection with M.marinum. In this study, we reported that VapA inhibiting macrophage autophagy by suppressing one of V-ATPase, ATPV0D1. We determined VapA interacted with ATPV0D1 through Pull-down, LC-MS/MS, and Co-IP analysis. Further, we reported that VapA affected macrophage loss of acidification of endo-lysosomal function by inhibiting the expression of ATP6V0D1.We also analyzed the effect of ATP6V0D1 on the intracellular proliferation of R.equi. We observed that the proliferation of R.equi was associated with the expression level of ATP6V0D1, when overexpressing ATP6V0D1 in macrophage increased the virulence R.equi103+ intracellular proliferation. The graphical diagram of this study is shown in Fig.6. In summary, we demonstrated that in some extend virulence *R.equi* escaped macrophage autophagy is caused lysosomal degradation mediated by VapA inhibiting ATP6V0d1.

## Funding

Please add: This research was funded by National Natural Science Foundation of China (31960694, 31760035), Excellent Youth Project of Ningxia Natural Science Foundation (2021AAC05008), Ningxia Overseas Students Innovation and Entrepreneurship Project.

## Institutional Review Board Statement

Not applicable

## Informed Consent Statement

Not applicable

## Conflicts of Interest

The authors declare no conflict of interest.

